# PTBP1 Regulates Injury Responses and Sensory Pathways in Adult Peripheral Neurons

**DOI:** 10.1101/2020.10.22.348631

**Authors:** Stefanie Alber, Pierluigi Di-Matteo, Matthew D. Zdradzinski, Letizia Marvaldi, Riki Kawaguchi, Katalin F. Medzihradszky, Ella Doron-Mandel, Nataliya Okladnikov, Ida Rishal, Reinat Nevo, Seung Joon Lee, Pabitra K. Sahoo, Alma L. Burlingame, Giovanni Coppola, Jeffery L. Twiss, Mike Fainzilber

## Abstract

Polypyrimidine Tract Binding Protein 1 (PTBP1) is expressed only at embryonic stages in central neurons. Its downregulation triggers neuronal differentiation in precursor and non-neuronal cells, an approach recently used to generate neurons *de novo* for amelioration of neurodegenerative disorders. Moreover, PTBP1 is replaced by its paralog PTBP2 in mature central neurons. Surprisingly, we found both proteins co-expressed in adult sensory and motor neurons, with PTBP2 restricted mainly to the nucleus, while PTBP1 shows strong axonal localization. Levels of axonal PTBP1 increased markedly after peripheral nerve injury, and its cargos include mRNAs involved in axonal growth and regeneration, such as importin β1 and RhoA. Perturbation of PTBP1 affects neuronal injury responses, axon outgrowth and sensation *in vivo*. Thus, PTBP1 has roles in sensory function and regenerative capacity of adult sensory neurons. These findings suggest that caution may be required before considering targeting PTBP1 for therapeutic purposes.

## Introduction

Axonal regeneration requires elongating growth from the proximal nerve segment after injury. Although functional regeneration occurs in the peripheral nervous system (PNS), central nervous system (CNS) neurons must contend with a lower intrinsic growth capacity as well as a growth inhibitory extracellular environment (Mahar and Cavalli, 2018). What are the intrinsic differences that enable more efficient regeneration in the PNS?

PNS regeneration requires activation of intrinsic growth programs by retrograde signaling from injured axons (Rishal and Fainzilber, 2014). An extensive series of studies have shown that intra-axonal protein synthesis is required for both retrograde injury signaling and for subsequent axonal regeneration (Smith et al., 2020). The RNA Binding Proteins (RBP) that transport regeneration associated mRNAs to axons are a critical regulatory node in the system (Turner-Bridger et al., 2020). Sensory axons contain a plethora of RBPs (Lee et al., 2018), however it is still unclear how dozens of RBPs associated with thousands of transcripts are localized and regulated to support axonal growth and maintenance. Misregulation of RBPs can cause serious damage in the nervous system and has been recognized as a critical determinant of neurological diseases (Cosker et al., 2016; Nussbacher et al., 2019).

The current study originated in a search for the RBPs required for axonal localization of the mRNA encoding importin β1, a critical regulatory factor for retrograde injury signaling (Hanz et al., 2003; Perry et al., 2012). Polypyrimidine tract binding protein 1 (PTBP1) was one of the most prominent candidates we identified, and this was particularly intriguing since previous studies have established that PTBP1 is absent from mature CNS neurons, where it is replaced by its paralog PTBP2 (Keppetipola et al., 2012). Both paralogs have partially redundant roles in pre-mRNA splicing (Vuong et al., 2016). PTBP1 is highly expressed in proliferating CNS progenitors during development, but is down-regulated upon differentiation and is absent in mature CNS neurons. This effect can be attributed to the action of the neuron-specific microRNA miR-124, which promotes neuronal differentiation by reducing PTBP1 levels, with concomitant increase in PTBP2 (Makeyev et al., 2007). Strikingly, we found both proteins coexpressed in adult sensory neurons. Levels of axonal PTBP1 increased markedly after peripheral nerve injury, and perturbation of PTBP1 affects neuronal injury responses, axon outgrowth and sensation *in vivo*. These findings suggest that there are fundamental differences in the roles of PTBP1 in CNS versus PNS neurons.

## Results

### PTBP1 is associated with KPNB1 mRNA in adult peripheral neurons

PTBP1 was identified by RNA affinity chromatography aimed to identify RBPs associated with the RNA localization motif of importin β1 (Perry et al., 2016) (KPNB1) in sciatic nerve axoplasm (Fig. 1A-C, Table S1) and direct interaction was confirmed *in vitro* (Fig. S1A). PTBP1 is a multifunctional RNA binding protein with high expression levels in many cell types, except for mature neurons where it is typically replaced by its neuronal paralog PTBP2 (Hu et al., 2018; Keppetipola et al., 2012; Lillevali et al., 2001; Makeyev et al., 2007). We found that PTBP1 and PTBP2 are co-expressed in adult dorsal root ganglia (Fig. 1D-G, Fig. S1B,C), a striking finding given the notion that PTBP1 is not expressed in adult neurons. Immunostaining on lumbar dorsal root ganglia (DRG) sections detected PTBP1 expression in both glial cells and neurons (Fig. 1E-G, Fig. S1D-G). PTBP1 protein was present in all DRG neurons, independent of their cell body or axonal area (Fig. S1D, E), and was also found colocalizing with *KPNB1* mRNA in sciatic nerve axons (Fig. 1H-J).

**Fig. 1:**
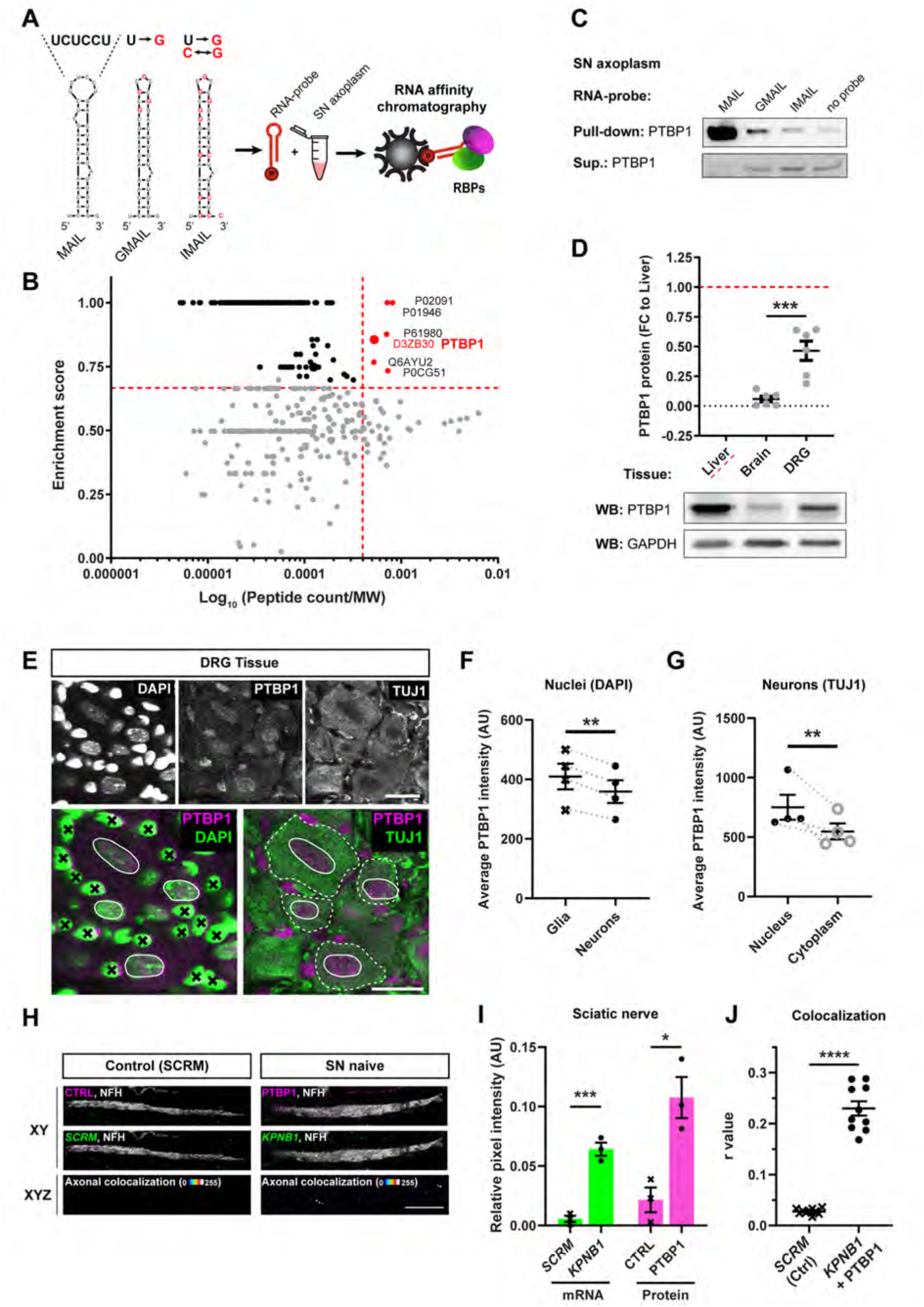
PTBP1 is expressed in adult sensory neurons and associated with *KPNB1* mRNA. (**A**) RNA structures and schematic of RNA affinity chromatography for mass spectrometry (MS) or western blot (WB). Secondary structures for MAIL (motif for axonal importin localization), and mutated GMAIL and IMAIL motifs, were generated using Mfold with modified bases shown in red (see Methods for sequences). (**B**) MS results for MAIL vs. GMAIL pulldown from rat axoplasm. Enrichment score (cutoff in red = 0.667) vs. Peptide count (PC)/ molecular weight (MW) shown in log scale (cutoff in red = 0.0004), for details see Methods. (**C**) WB for PTBP1 from mouse sciatic nerve (SN) axoplasm after pull-down with MAIL or control RNA motifs. Note that PTBP1 is enriched in the MAIL pulldown, while depleted from the supernatant. Similar results were obtained from rat and bovine axoplasm (data not shown). (**D**) WB analysis for PTBP1 protein levels in liver, brain and dorsal root ganglia (DRG) from adult mice. PTBP1 levels were normalized to GAPDH and shown as fold change (FC) from liver. Individual datapoints with mean ± SEM, unpaired t-test for Brain vs. DRG, *** indicates p< 0.001, n = 6 mice. (**E**) Tissue sections of adult lumbar DRG ganglia, stained for DAPI, PTBP1 and TUJ1 (Tubulin Beta 3 Class III). Scale bar = 20 μm. Merged images show outlined neuronal nuclei (white line) and cytoplasm (white dashed line), glia nuclei are marked by a black cross. (**F**) Quantification of PTBP1 signal in nuclei shows PTBP1 expression in both cell types with enrichment in glia. Individual datapoints with mean ± SEM, ratio paired t-test, ** indicates p< 0.01, n = 4 mice (> 100 nuclei for each mouse). (**G**) Quantification of PTBP1 in Tuj1 positive neurons in nucleus and cytoplasm. Individual datapoints with mean ± SEM, ratio paired t-test, ** indicates p< 0.01, n = 4 mice (> 60 cells for each mouse). Grey dashed lines indicate paired datapoints. **H-J**, Longitudinal sections of SN stained for NFH (Neurofilament heavy chain) and PTBP1 (protein), combined with fluorescent *in situ* hybridization (FISH) for *KPNB1* mRNA. No primary antibody was used as control for PTBP1 staining (CTRL) and *scramble (SCRM)* mRNA for *KPNB1* FISH. (**H**) Representative images of the staining (top), FISH (middle) and axonal colocalization of PTBP1 with *KPNB1* or *SCRM* mRNA in white (bottom). Scale bar = 10 μm. (**I**) Quantification of *KPNB1* mRNA vs. *SCRM* and PTBP1 protein vs. CTRL in SN. Separate quantification for mRNA and protein signal, individual datapoints with mean ± SEM, t-test, * indicates p< 0.05, *** indicates p< 0.001, n = 3 mice. (**J**) Quantification of axonal colocalization by Person’s correlation. R-values as individual datapoints with mean ± SEM, t-test, **** indicates p< 0.0001, n = 10 sciatic nerve sections from 3 mice.

### Axonal PTBP1 is elevated after nerve injury

As *KPNB1* and other axonal mRNAs are involved in injury responses, we monitored PTBP1 levels in sciatic nerve axons over time after lesion. PTBP1 levels start to increase in axoplasm three days after injury, reaching an order of magnitude elevation after one week (Fig. 2A).

**Fig. 2.**
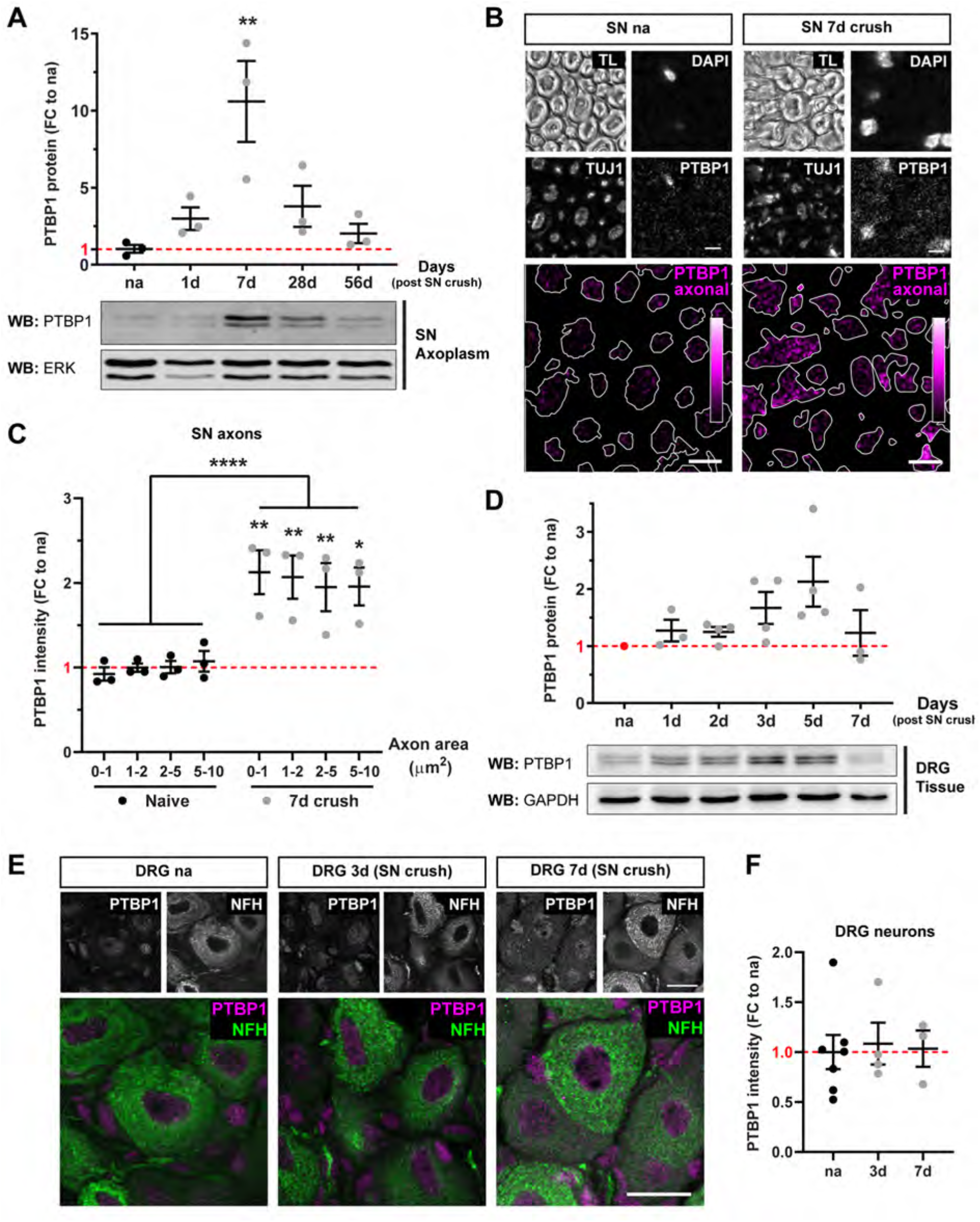
Increased PTBP1 in axons after nerve lesion. **(A)** PTBP1 protein expression in sciatic nerve (SN) axoplasm under naive (na) and injury (inj) conditions. Detection by western blot (WB). PTBP1 levels were normalized to Extracellular Signal-Regulated Kinase 1/2 (ERK) and the data is shown as fold change (FC) to naive (red dashed line). Individual datapoints with mean ± SEM, one-way ANOVA with Tukey’s multiple comparison test, ** indicates p< 0.01, n = 3 mice for each timepoint. **(B)** Representative images of sciatic nerve cross sections from naive nerve and 7d after crush injury. Axonal PTBP1 is shown in magenta hot scale (0-255), outline of TUJ1 (Tubulin Beta 3 Class III) indicated with a white line, TL (transmission light), scale bar = 5 μm. **(C)** Quantification of axonal PTBP1 signal in TUJ1 positive axons of different sizes (Binning of axon area: 0-1, 1-2, 2-5, 5-10 μm^2^). PTBP1 intensity is shown as FC to naive (red dashed line), individual datapoints with mean ± SEM, two-way repeated measure (RM) ANOVA, **** indicates p< 0.0001, Sidak’s multiple comparison test, ** indicates p< 0.01, * indicates p< 0.05, n = 3 (>700 axons/mouse for each size category). **(D)** PTBP1 protein expression in cell bodies of dorsal root ganglia (DRG). WB of PTBP1 with normalization to GAPDH (Glyceraldehyde-3-Phosphate Dehydrogenase), shown as FC to naive (red dashed line). Individual datapoints with mean ± SEM, one-way ANOVA with Tukey’s multiple comparison test, not significant, n = 3-4 mice for each timepoint. **(E)** Representative images of DRG tissue sections from naive, 3d and 7d after SN crush. Scale bar = 20 μm. **(F)** Quantification of average PTBP1 signal in DRG neurons (summary of all cell sizes, NHF+ & – cells). Individual datapoints with mean ± SEM, one-way ANOVA with Tukey’s multiple comparison test, not significant, n = 3-7 mice (> 95 cells/mouse).

PTBP1 was maintained at elevated levels for up to 21 days, then declined back to baseline levels after 2 months (Fig. S2A). Robust upregulation was observed within sensory axons of all diameters and subtypes at the one week time point (Fig. 2B, C, Fig. S2B, C). Motor axons also contained PTBP1, with more variable injury-induced upregulation (Fig. S2D), and glial PTBP1 levels per cell were not changed after injury (Fig. S2E). Interestingly there was also little or no increase in PTBP1 in DRG neuron cell bodies after sciatic nerve injury (Fig. 2D-F, Fig. S2F-H), indicating that the vast majority of increased PTBP1 is transported to axons.

### PTBP1 traffics RNAs to regenerating axons

Transport of PTBP1 to axons might indicate that it functions as a carrier RBP to traffic RNAs required for neuronal regeneration. Hence, we carried out RNA sequencing on PTBP1 immuno-precipitates from sciatic nerve axoplasm before and after injury (Fig. 3A-D, Fig. S3A-C, Table S2). We identified 1069 mRNAs specifically associated with PTBP1 in axoplasm, 104 of which revealed similar enrichment in naive and injury conditions, 721 were highly enriched in naive nerves, and 244 mRNAs were enriched after injury (Fig. 3D). Ingenuity pathway analysis highlighted a plethora of signaling networks enriched after injury, suggesting that PTBP1-associated RNAs encode proteins involved in cytoskeleton remodeling and axon growth after nerve injury. RhoA, a well-characterized nerve regeneration regulator (Kalpachidou et al., 2019), was identified as one of the prominent PTBP1-associated mRNAs (Table S2).

**Fig. 3:**
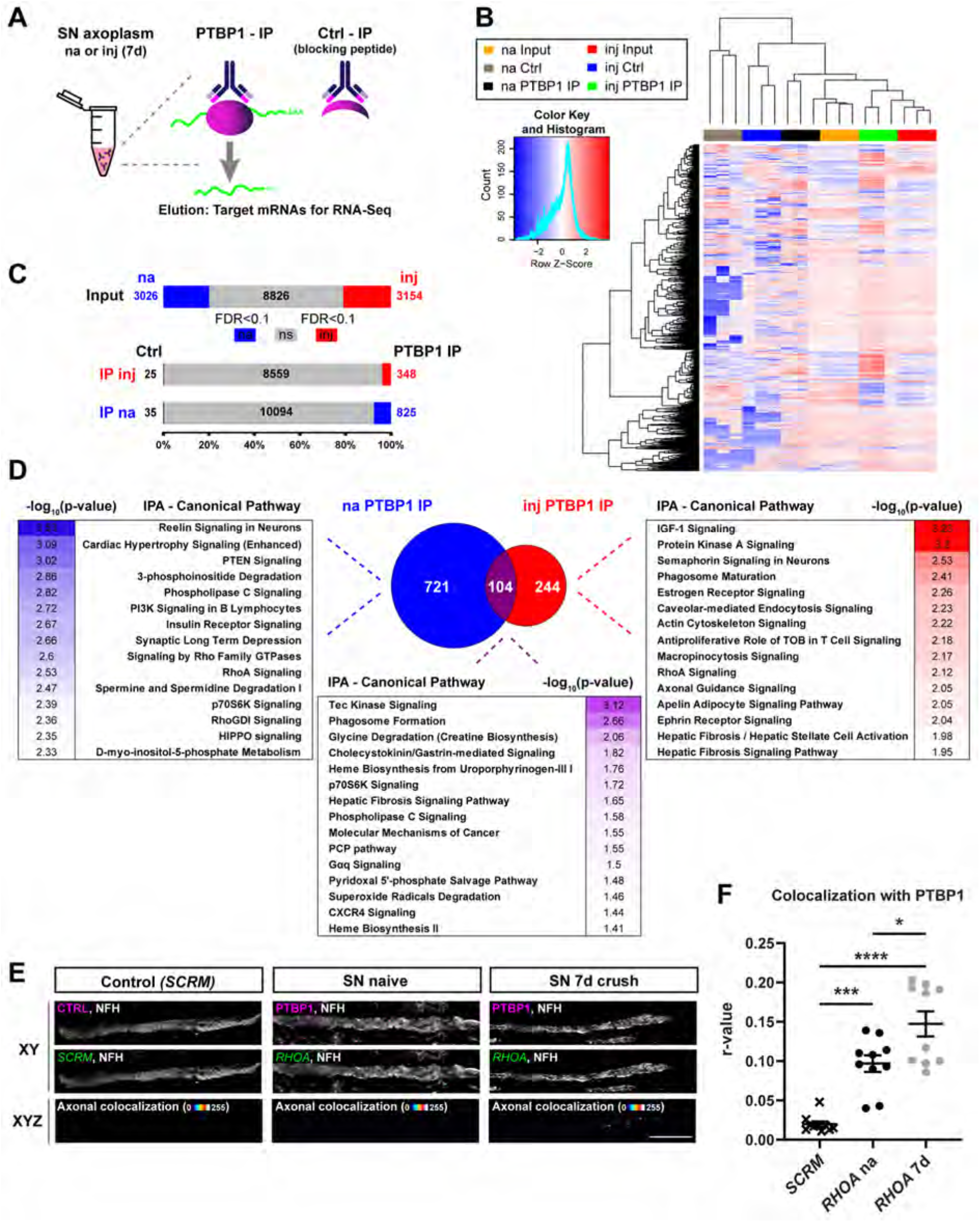
Identification of PTBP1 cargo mRNAs in axons. **(A)** Schematic of RNA coimmunoprecipitation (RNA-IP) with PTBP1 from naive (na) sciatic nerve (SN) axoplasm or 7d after crush injury (inj). A blocking peptide specific for the PTBP1 antibody was used as a negative control (Ctrl). **(B)** Heat map of differentially expressed genes. Top 2000 transcripts based on rank (FDR < 0.1) are shown with three replicates for each group. **(C)** Bar plot showing the total number of transcripts identified as differentially and non-differentially expressed (FDR<0.1). Grey = not significant (ns), blue = enriched in na, red = enriched in inj. **(D)** Comparison of PTBP1 binding transcripts between naive and injury. Venn diagram shows the number of unique or overlapping transcripts in naive/injury (enrichment criteria FDR < 0.1). Naive = blue, injury = red and overlap = purple. Side panels show top 15 canonical pathways enriched in each section (Ingenuity Pathway Analysis (IPA), Qiagen). Heat map shows – log10 (p-value) from minimum to maximum value (spectrum from white to saturated color for each section). **(E)** Representative images for RNA-FISH (Fluorescence in situ hybridization) for *RHOA* (Ras Homolog Family Member A) mRNA and PTBP1 protein in longitudinal sections of SN. Scale bar =10 μm. **(F)** RNA-FISH quantification of axonal colocalized PTBP1 and RHOA mRNA. Pearson correlation analysis, r-value of individual datapoints with mean ± SEM, one-way ANOVA with Tukey’s multiple comparison test, * indicates p< 0.05, *** indicates p< 0.001, **** indicates p< 0.0001, n = 10 nerve sections from 3 different mice.

We combined RNA fluorescent in situ hybridization (FISH) for *RhoA* with immunostaining for PTBP1 in sciatic nerve sections (Fig. 3E, F and Fig. S3D-G). There was significant colocalization between the two in both naive and injury conditions. However, overall levels of RhoA mRNA were extremely low in naive sciatic nerve and during the first few hours after crush (Fig. S3D), while increased association of RhoA mRNA with PTBP1 was observed seven days after crush injury (Fig. 3F), consistent with a role for PTBP1 in trafficking RhoA mRNA to regenerating axons.

### PTBP1 regulates injury responses and regeneration of adult nociceptive neurons

Is PTBP1 a critical factor for regeneration of adult sensory neurons? To address this question, we used AAV-encoded shRNA for knockdown of PTBP1 in DRG *in vivo* (Fig. 4A, B; Fig. S4A, B). Intrathecal injections of AAV9 served to deliver shPTBP1 or shCtrl, three weeks prior to a conditioning sciatic nerve lesion. Seven days after the lesion, we harvested sciatic nerve for immunohistochemical monitoring of regeneration, and L4/L5 lumbar DRG for neuronal cultures (Fig. 4C-G). PTBP1 knockdown greatly impaired the enhanced neuronal outgrowth typically caused by a conditioning lesion, with significant reductions observed both in the percentage of growing neurons and total neurite length (Fig. 4C-E). Moreover, injury-induced axonal expression of calcitonin gene-related peptide (CGRP) (Toth et al., 2009) was not detected in animals treated with shPTBP1 (Fig. 4F, G; Fig. S4C-E).

**Fig. 4:**
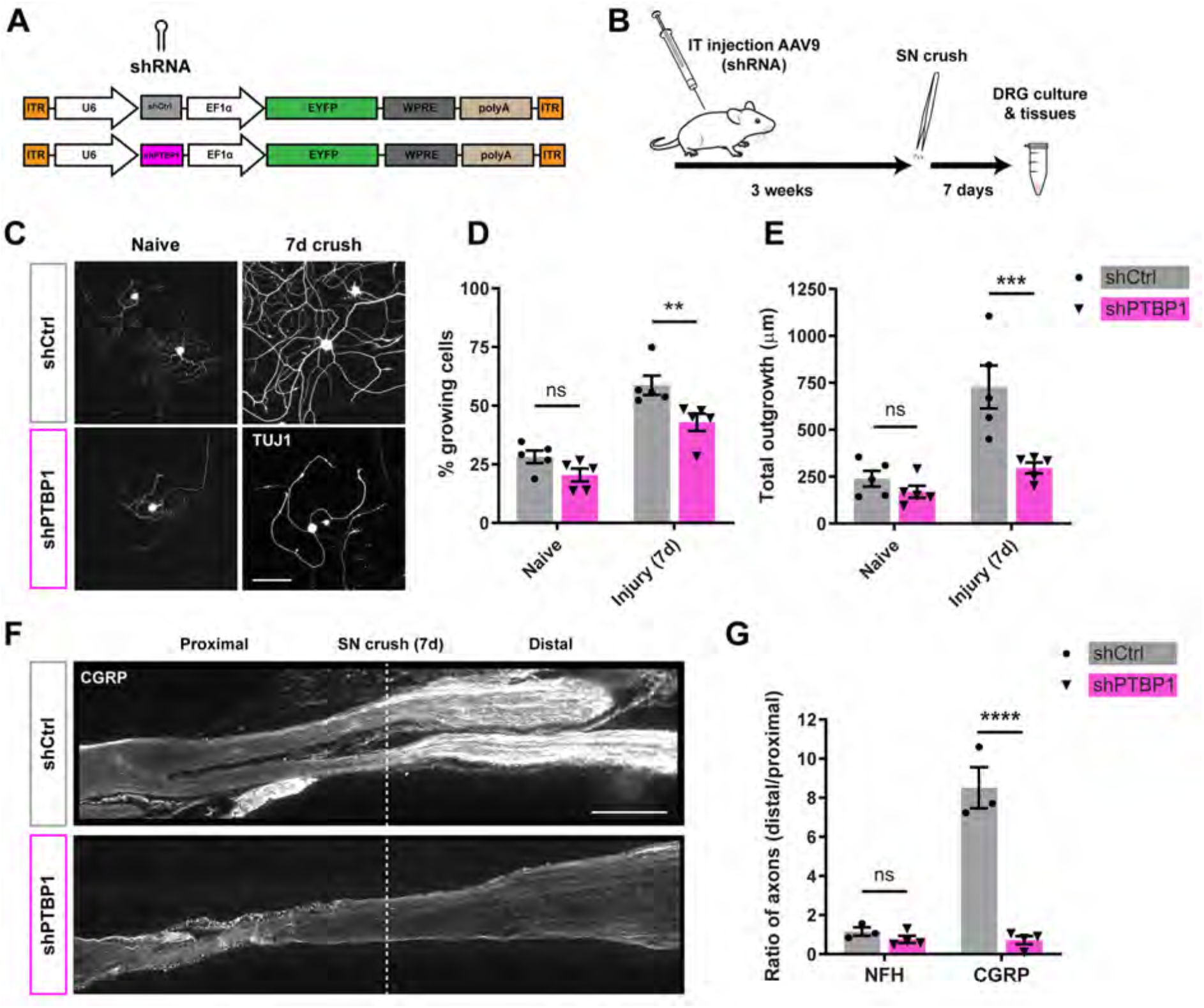
PTBP1 in the injury response of adult sensory neurons. (**A**) Viral constructs used for knockdown experiments. ITR (inverted terminal repeat); U6 (type III RNA polymerase III promoter), shRNA (small hairpin RNA), EF1α (elongation factor 1 alpha) promoter, EYFP (enhanced yellow fluorescent protein), WPRE (Woodchuck hepatitis virus posttranscriptional regulatory element), polyA (polyadenylation signal). (**B**) Schematic of *in vivo* knockdown experiments. shRNA constructs were delivered via AAV9 (Adeno associated virus serotype 9) through intrathecal (IT) injection, followed by sciatic nerve (SN) crush 3 weeks later. At day 7 after crush DRG cells were collected for culture and SN for immunohistochemistry. (**C**) Representative images for DRG neurons in culture (20 h after plating) for naive and injury conditioned cells from shCtrl and shPTBP1 treated mice. TUJ1 (Tubulin Beta 3 Class III), scale bar = 100 μm. **D** & **E**, Quantification of growing cells in **(c)**. Individual datapoints with mean ± SEM, two-way RM (repeated measure) ANOVA with Sidak’s multiple comparison test, ** indicates p< 0.01, *** indicates p< 0.001, n = 5 mice per condition. (**D**) Percent of growing cells. (**E**) Total outgrowth in μm. (**F**) SN longitudinal sections 7 days after crush for shCtrl and shPTBP1. Staining for CGRP (Calcitonin Gene-Related Peptide 1), injury site indicated by the white dashed line, scale bar = 500 μm. (**G**) Quantification of the number of positive axons (average of several regions up to 4 mm) proximal and distal to the crush site, shown as ratio distal/proximal for NFH (Neurofilament heavy chain, images in Fig. S4C) and CGRP. Individual datapoints with mean ± SEM, twoway RM ANOVA with Sidak’s multiple comparison test, p-value **** < 0.001, n = 3-4 mice.

We next used behavioral assays to assess the functional effects of PTBP1 depletion in the injury paradigm (Fig. 5A). Knockdown of PTBP1 had no effect on naive or injury-associated gait parameters assessed on a CatWalk apparatus (Fig. 5B, Fig. S5A), but interestingly partly ameliorated the reduced mechanical and thermal sensitivity observed in the injured limb seven days post-lesion (Fig. 5C, D, Fig. S5B, C). Independent testing of these findings was then carried out using a decoy RNA oligonucleotide that transiently blocks PTBP1 (Denichenko et al., 2019). PTBP1-blocking or control oligonucleotides were injected locally to the sciatic nerve just before lesioning (Fig. 5E). This experimental paradigm enables local perturbation of PTBP1 in axons, as the decoy oligo does not reach neuronal cell bodies (Fig. S5D-F). Effects of the PTBP1 blocking oligonucleotide in axons were similar to those observed previously by shRNA knockdown, namely no effects on gait parameters (Fig. 5F) and amelioration of the injury-induced changes in mechano- and thermosensitivity (Fig. 5G, H; Fig. S5G-I). Taken together, these data indicate that axonal PTBP1 regulates injury responses and regeneration of adult nociceptive neurons.

**Fig. 5:**
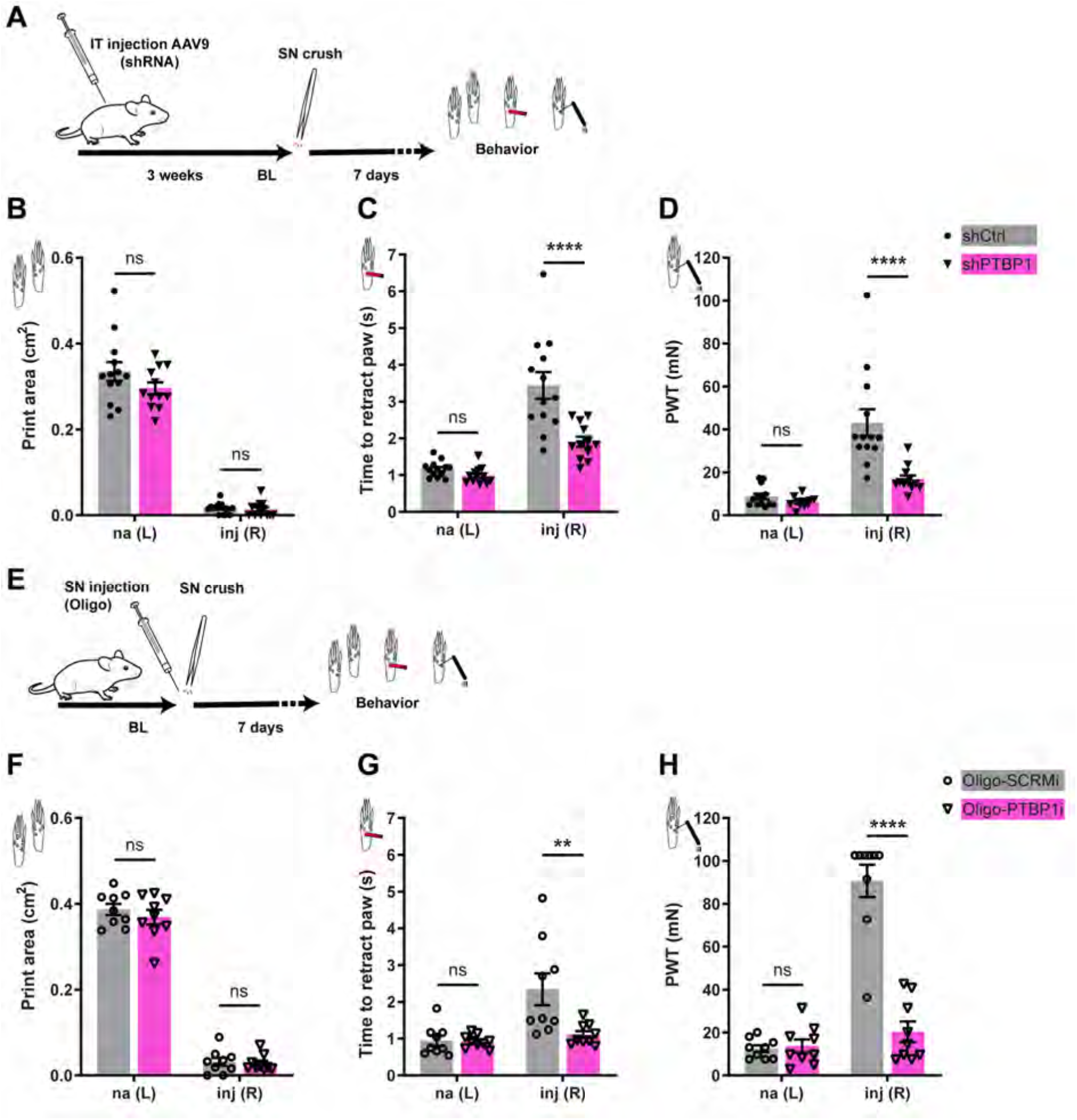
Functional consequences of PTBP1 perturbation after nerve crush. **(A)** Timeline for behavioral assays after knockdown of PTBP1 in DRG neurons. shRNA constructs were delivered via AAV9 (Adeno associated virus serotype 9) through intrathecal (IT) injection. Baseline (BL) testing was performed 3 weeks later, just prior to sciatic nerve (SN) crush. At day 7 after crush, both hindlimbs (naive left (na L), right injury (inj R) were tested in CatWalk gait analysis **(B)**, heat probe **(C)** and von Frey mechanosensation **(D)** paradigms. B-D, Behavior after knockdown of PTBP1 7 days after SN crush. Individual datapoints with mean ± SEM, two-way repeated measure (RM) ANOVA with Sidak’s multiple comparison test, **** indicates p< 0.0001, n = 11-13 mice. PWT (paw withdrawal threshold). **(E)** Timeline for behavioral assays after inhibition of PTBP1 in the SN. BL was established prior to injections/crush. Inhibitory decoy RNA oligos (PTBP1i or SCRMi (scramble)) were injected directly into the SN, followed immediately by SN crush. After 7 days, mice were tested in CatWalk gait analysis **(F)**, heat probe **(G)** and von Frey paradigms **(H)**. F-H, Behavior after inhibition of PTBP1 7 days after SN crush. Individual datapoints with mean ± SEM, two-way RM ANOVA with Sidak’s multiple comparison test, ** indicates p< 0.01, **** indicates p< 0.0001, n = 9 mice.

## Discussion

The findings that PTBP1 is expressed in adult sensory and motor neurons, and that it has functional roles in sensation and regeneration in adult neurons, are surprising in light of the well-established model that PTBP1 downregulation triggers neuronal differentiation, and it is not expressed in adult CNS neurons (Hu et al., 2018; Vuong et al., 2016). A previous bioinformatics study reported that mature sensory neurons have splicing profiles consistent with continued expression of PTBP1 in adulthood (Weyn-Vanhentenryck et al., 2018), but did not provide experimental evidence to confirm that prediction. Our study provides experimental data on multiple levels for PTBP1 expression in adult sensory neurons, and moreover highlights a new role for PTBP1 as an axonal carrier of mRNAs required for growth and regeneration.

A number of groups recently took advantage of the canonical roles of PTBP1 in splicing in early development and in driving neuronal differentiation (Xue et al., 2013; Xue et al., 2016) to assess the feasibility of in vivo differentiation of new neurons to ameliorate neurodegeneration. Zhou and colleagues used viral delivery of a PTBP1 gene editing system to convert Muller glia into retinal ganglion cells, alleviating symptoms of RGC loss (Zhou et al., 2020). They further reported that this approach could induce dopaminergic neurons in the striatum to alleviate Parkinson’s disease deficits. Another study also targeted Parkinson’s disease by conversion of astrocytes to neurons by depleting PTBP1 (Qian et al., 2020). These studies are promising proofs of concept for treating neurodegeneration by targeting PTBP1 (Arenas, 2020), but they are based on the notion that PTBP1 is not expressed and does not have functions in adult neurons. Our current work clearly shows that is not the case for peripheral sensory and motor neurons, suggesting caution will be required in order to advance PTBP1 targeting approaches to the clinic.

## Acknowledgements

We are thankful to Vladimir Kiss and Mathias Mahn for help and advice on imaging, Michael Tsoory for guidance with behavioral assays, and Dalia Gordon and Rotem Perry for helpful discussions. We gratefully acknowledge funding from the Dr. Miriam and Sheldon G. Adelson Medical Research Foundation (A.L.B., G.C., J.L.T., and M.F.), the International Foundation for Research in Paraplegia (IRP Grant # P177 to M.F.), the National Institutes of Health (R01-NS117821 to JLT, P41GM103481 and 1S10OD016229 to ALB) and the South Carolina Spinal Cord Injury Research Fund (2019-PD-02 to PKS). M.F. is the incumbent of the Chaya Professorial Chair in Molecular Neuroscience, and J.L.T. is the incumbent of the Smartstate Chair in Childhood Neurotherapeutics.

## Author Contributions

S.A. and M.F. designed the study, S.A., P.D.M., M.D.Z., L.M., E.D.M., N.O., and P.K.S. conducted experiments, M.D.Z. and P.K.S. carried out FISH analyses, R.K. performed RNA-seq, K.F.M. performed mass spectrometry, R.N. conducted imaging analyses, I.R., A.L.B., G.C., J.L.T., and M.F. supervised research, S.A. and M.F. wrote the manuscript draft with input from all the co-authors.

## Supplementary Material

### Animals

All animal procedures were performed in accordance with the guidelines of the Weizmann Institute of Science and University of South Carolina (USC) Institutional Animal Care and Use Committees (IACUCs). Adult C57BL6 and Hsd:ICR mice were purchased from Envigo Ltd (Israel) and maintained at the veterinary resources department of the Weizmann Institute with free access to food and water. Wistar rats (Envigo) were used for mass spectrometry analyses. All experiments were performed on male animals between 2-5 months of age.

### Sciatic nerve crush model

Animals were anesthetized with ketamine/xylazine (10 mg/kg body weight intraperitoneal). Sciatic nerve (SN) crush was performed at mid-thigh level using a fine jewelers’ forceps in 2 adjacent positions for 30 seconds each. Only the right side was subjected to the crush injury procedure (inj), the contralateral side served as uninjured, naive (na) control.

### Behavior

All mice that were used for behavioral testing were kept on a reverse dark/light cycle so that the assays could be performed during the “dark” active phase of the animal. Before each session a 1 h habituation period was given in the test room. All behavioral tests were done in blind.

#### CatWalk

gait analysis was performed using CatWalk XT setup and software (CatWalk XT 10.6, Noldus Information Technology, The Netherlands) as previously described (Perry et al., 2012). This setup allows for quantification of footprints and gait in unforced moving animals. Motivation was achieved by placing the home cage at the end of the runway. At least 3 compliant runs per animal were recorded in each session.

#### Heat probe

This test was used to evaluate the response to noxious heat, as previously described (Marvaldi et al., 2020). While holding the animal, a metal probe, heated to 58 °C, was applied to both hind paws consecutively (middle part of the plantar side, left paw naive, right paw after crush injury). The paw withdrawal latency was recorded in seconds (s) for 3 trials, with 20-minute intervals between repeats.

#### von Frey

Mechanical sensitivity was tested in the von Frey paradigm as previously described (Marvaldi et al., 2020). Mice were placed in an elevated setup of transparent chambers with a wire mesh grid on the bottom. Following a 1 h habituation period, nylon monofilaments of different diameter (von Frey Hairs #37450-275, ugo basile) were pressed against the plantar surface of the hind paw until bending of the filament and held for a maximum of 3 s. A positive response was noted if the paw was sharply withdrawn upon application of the filament. The test was started with a stimulus of 1.4 g (filament target force of 13.7 millinewtons (mN)) and continued in an up-down testing paradigm for at least 5 representations of the stimulus. The paw withdrawal threshold (PWT) was calculated for each animal using UDMAP (V3.0).

### Reagents and antibodies

Primary antibodies (ABs) used in this study for immunoprecipitation (IP), immunocytochemistry (ICC), immunohistochemistry (IHC), immunofluorescence (IF: ICC & IHC) and western blot (WB) were: Anti-PTBP1 (Santa Cruz Biotechnology, sc-16547, used for IP 2.5 μg/25 μl beads, IF 1:200 and WB 1:500, used for all main experiments until it was discontinued in 2016), anti-PTBP1 (abcam, ab5642, ICC 1:500, IHC 1:200, WB 1:1000; used as replacement for sc-16547 as it recognizes the same epitope). The blocking peptide for PTBP1 antibodies (sc-16547 and ab5642) was synthesized by Sigma ([H]-DGIVPDIAVGTKRGSDELFS-[OH]). Furthermore, anti-PTBP1 (abcam, ab133734, WB 1:1000), anti-PTBP2 (Abnova, H00058155-A01, IF 1:1000, WB 1:2000), anti-PTBP2 (abcam, ab154787, WB 1:1000). anti-ROD1 (Santa Cruz, sc-100845, WB 1:200), anti-GFP (abcam, ab6556, IHC 1:500, ICC & WB 1:5000), anti-NFH (abcam, ab72996, IHC 1:1000, ICC 1:2000), anti-GAPDH (Santa Cruz Biotechnology, sc-32233, WB 1:5000), anti-ERK1/2 (Sigma, M5670, WB 1:30000), anti-β-III tubulin TUJ1 (abcam, ab18207, IF 1:1000, WB: 1:6000), anti-CGRP (AbD SEROTEC 1720-9007, IF 1:1000), anti-Nucleolin (abcam, ab50279, IF & WB 1:1000). DAPI (Thermo Fisher Scientific, D1306, 1μg/ml).

Secondary antibodies used for IF are donkey anti-mouse/rabbit/goat/chicken Alexa Fluor 488/594/647 (Jackson ImmunoResearch, IHC 1:500, ICC 1:1000). Secondary HRP-conjugated mouse and rabbit antibodies for Western blot were purchased from Bio-Rad Laboratories, secondary bovine anti-goat IgG-HRP (Santa Cruz Biotechnology, sc-2350). All HRP-conjugated antibodies were used 1:10000.

### Protein extraction and SDS-PAGE

For total protein extraction, tissues (liver, brain, DRG) were collected in RIPA buffer (150 mM NaCl, 1.0 % NP-40, 0.5 % sodium deoxycholate, 0.1 % SDS, 50 mM Tris, pH 8.0.), supplemented with protease/phosphatase/RNase inhibitors. Tissue samples were homogenized in Dounce Tissue Grinders (WHEATON 33, 1 ml #357538, 7 ml #357542) or, using plastic pestles in an Eppendorf tube.

To obtain neuron enriched protein extracts from DRG tissue, ganglia were dissociated according to the DRG culture protocol described below. After the percoll-gradient (depletion of non-neuronal cells) cell pellet was washed in PBS and then lysed in RIPA buffer.

Axoplasm for biochemical analysis was extracted from mouse or rat SN as previously described (Doron-Mandel et al., 2016). To minimize glia contamination, transport buffer (20 mM HEPES, 110 mM KAc, 5 mM MgAc, pH 7.4, supplemented with protease/phosphatase/RNase inhibitors) was used in this extraction protocol.

All protein extracts were incubated on ice for 20 min, followed by a spin down at 10000 g for 10 min at 4 °C. Supernatant was used for protein electrophoresis (Mini-PROTEAN^®^ Tetra Cell Systems, Bio-Rad) and blotting (Trans-Blot^®^ Turbo™ Transfer System (Bio-Rad). Chemiluminescence was detected with ImageQuant LAS 4000 (GE Healthcare) and band intensities were quantified using built-in software.

### Pull-down of RNA binding proteins

Biotinylated ribonucleic acid (RNA) probes for pull-down assays were synthesized by IDT (Integrated DNA Technologies, Syntezza, Israel):

MAIL: 5’-Biotin-TEG-UCACAAACAAGCUCUCUCCUGACUUGUAUUGUGG-3’;

GMAIL: 5’-Biotin-TEG-UCACAAACAAGCGCGCGCCGGACUUGUAUUGUGG-3’;

IMAIL: 5’-Biotin-TEG-UGAGAAAGAAGGGCGCGCCGGACUUCUAUUCUCC-3’;

ZIP (ACTB): 5’-Biotin-TEG-ACCGGACUGUUACCAACACCCACACCCCUGUGAUGAAACAAAACCCAUAAAUGC-3’ (Kim et al., 2015). RNA affinity chromatography was performed as previously described (Doron-Mandel et al., 2016). Briefly, streptavidin magnetic beads were washed several times and incubated with the biotinylated probe for 1 h at 4 °C, followed by 3 additional washes. Meanwhile, freshly prepared axoplasm was applied to beads without a probe for 1 h to deplete unspecific proteins. Afterwards the unbound fraction was added to the specific probes for 1 h at 4 °C.

For the *in vitro* binding assay, purified recombinant human PTBP1 protein was purchased from Abnova (H00005725-P01). 1 μg of recombinant PTBP1 was used for each probe (MAIL, IMAIL, ZIP with 20 μl streptavidin beads).

Bound material was washed and eluted from the beads using SDS sample buffer (western blot, WB) or RNaseA (for mass spectrometry, MS). The samples were loaded onto 10 % SDS-PAGE gels, followed by WB or MS analysis.

### RNA-binding protein identification using mass spectrometry

#### Sample preparation for mass spectrometry

Importin-beta1-mRNA-loop-binding proteins and proteins isolated using a control RNA-loop were fractionated on SDS-PAGE (Biorad tris-glycine precast gels 4-15%), and stained using Colloidal Blue Staining Kit (Invitrogen). Each lane was cut into 10 pieces and these pieces were subjected to our in-gel digestion protocol [http://ms-facility.ucsf.edu/protocols.html]. Briefly, the SDS and CBB were removed using 25 mM ammonium bicarbonate (ABC) in 50% acetonitrile/water, then the disulfide bridges were reduced with DTT in 25 mM ABC buffer, and the free –SH groups were alkylated with iodoacetamide. Later the gels were dehydrated, and reconstituted with the trypsin solution in 25 mM ABC buffer. 4h digestion was performed, at 37·C, the resulting peptides were extracted with 5% formic acid in 50% acetonitrile/water. The peptide solutions were concentrated and submitted to LC/MS/MS analysis.

#### Mass spectrometry analysis

The LC/MS/MS analysis was performed using a NanoACQUITY-LTQ-Orbitrap XL sytem (Waters, Thermo Scientific, respectively). The peptide fractionation was performed on a C18 column (75μm x 150 mm) at a flowrate of ~400 μl; solvent A was 0.1% formic acid in water, solvent B was 0.1% formic acid in acetonitrile. A linear gradient was applied from 2% to 35% organic in 40 minutes. Data were acquired for 60 min following sample injection, in a data-dependent manner. The precursor ions were measured in the Orbitrap, and the 6 most abundant multiply charged ions were selected for collision-induced dissociation experiments performed and measured in the linear trap. The trigger threshold was 1000, and dynamic exclusion was enabled.

#### Data processing

An in-house script, PAVA was used for peak-picking, and the database searches were performed with the 10 peaklists representing the same sample combined, using Protein Prospector. <u>Search parameters</u>: only tryptic peptides were permitted, with 2 missed cleavages; mass accuracy within 20 ppm and 0.6 Da for precursors and fragments, respectively; fixed modification: Cys carbamidomethylation; variable modifications: Met(O); protein N-acetylation; N-terminal Gln->pyroGlu. The database was UniProtKB.2013.6.17, concatenated with random sequences for each entry, *Rattus* and *Mus musculus* proteins were selected, human keratins were added to the list (108456 entries were searched). <u>Acceptance criteria</u>: minimum score: 20 & 15; max E-value: 0.05 & 0.1 for proteins and peptides, respectively. FDR < 1%, based on the number of decoy hits.

Peptide counts (PC) have been normalized by the molecular weight (MW) of the identified protein. Enrichment of MAIL vs. GMAIL (control) has been calculated as:

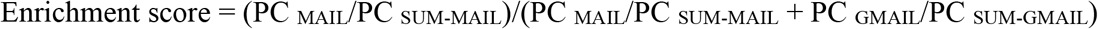

Proteins (peptides) only identified in the MAIL pulldown have an enrichment score of 1, whereas peptides only identified in GMAIL control have an enrichment score of 0. To identify potential candidates, a 2-fold enrichment of MAIL vs. GMAIL was used (Enrichment score > 0.6667), as well as a cutoff for high coverage with PC/MW > 0.0004. A complete list of identified proteins can be found in Table S1.

### DRG culture

The procedure for DRG neuron culture was performed as previously described (Rishal et al., 2012). Briefly, adult mouse DRGs were dissociated with 100 U of papain (P4762, Sigma) followed by 1 mg/ml collagenase-II (11179179001, Roche) and 1.2 mg/ml dispase-II (04942078001, Roche). The ganglia were then triturated in HBSS, 10 mM Glucose, and 5 mM HEPES (pH 7.35) using a fire coated Pasteur pipette. Neurons were layered on 20 % percoll in L15 media and recovered through centrifugation at 1000 g for 8 min. Cells were washed briefly in growth media (F12, 10 % Fetal Bovine Serum (FBS), Primocin (100μg/ml, InvivoGen #ant-pm-1)) and plated on poly-L-lysine (P4832, Sigma) and laminin (23017-015, Invitrogen) coated glass cover slips. Culture media and serum from Thermo Fisher Scientific.

### Immunocytochemistry (ICC)

Cultured cells were washed in PBS and fixed in 4 % PFA for 20 min. Blocking was performed in 10 % donkey serum, 1 mg/ml bovine serum albumin (BSA), 0.2 % Triton in PBS for 1 h at room temperature (RT, ~25°C). Primary antibodies were incubated in AB-solution (5 % donkey serum, 1 mg/ml BSA, in PBS) for 1 h at RT or overnight at 4 °C. Secondary ABs (raised in donkey, 1:1000, Jackson ImmunoResearch) were applied in AB-solution for 2 h at RT. For imaging, coverslips were mounted with Fluoromount™ Aqueous Mounting Medium (F4680, Sigma Aldrich).

### Histology, immunofluorescence (IF) and image analysis

Tissues (DRGs or sciatic nerves) were fixed in 4 % PFA for 24h at 4 °C, then washed in PBS, followed by incubation in 30 % sucrose for 48 hours at 4 °C. Tissues were embedded in O.C.T. Compound (Scigen, 4583) and 10 μm sections were obtained (cross-sections for sciatic nerve). Blocking was performed in 10 % horse serum, 0.2 % Triton X-100 in PBS for 2 h at RT. Primary antibodies were incubated in AB-solution (5% horse serum, 0.2 % Triton X-100 in PBS) overnight at 4 °C. Secondary ABs were applied in AB-solution for 2 h at RT. Slides were mounted in Fluoromount (F4680, Sigma Aldrich) and confocal images were taken in the following days. Samples were imaged with a DMi8 Leica (Leica Microsystems, Mannheim, Germany) confocal laser-scanning microscope, using a HC PL APO 40x/1.3 oil-immersion objective and HyD SP GaAsP detectors. Imagestacks of 4 μm were collected for each sample, with 0.5 μm thick optical sections. Images were acquired by maintaining a pixel size of 0.142 μm and an image dimension of 2048 × 2048 pixels.

### Quantification of PTBP1 levels in DRG sections

For each mouse, L4 and L5 ganglia were quantified from both sides (left naive, right after SN crush injury) and 8-10 non-overlapping sections were imaged and used for analysis. For the automated analysis on neuronal or glial nuclei, positive objects were distinguished based on DAPI intensity signal, size and morphology, using Ilastik 1.0 (Berg et al., 2019). PTBP1 levels were also evaluated using a customized protocol. For this analysis, neuronal cell bodies were outlined according to TUJ1 or NFH staining. Only neurons showing a clear nuclear signal (DAPI) were selected for quantification. Measurements of average PTBP1 intensity were obtained using ImageJ/Fiji (Schindelin et al., 2012). DAPI was used to identify the nuclear compartment and was subtracted from the total cell body area to obtain the cytoplasmic compartment. For analysis based on cell size, the cell body area (μm^2^) was quantified and neurons were divided in size bins: 0-200, 200-300, 300-500, >500 μm^2^.

### Quantification of PTBP1 levels in sciatic nerve sections

Per each mouse, eight images (each one containing hundreds of axons) were acquired from non-consecutive sections, from naive or crushed nerve (1mm proximal to the injury site). Axons were identified using different markers: TUJ1, NFH or ChAT. Measurements of average PTBP1 intensity were obtained using ImageJ/Fiji, only the intensity of PTBP1 inside the axons was considered for the analysis. For analysis based on axon size, the axon area (μm^2^) was measured and neurons were divided in size bins: 0-1, 1-2, 2-5, 5-10 μm^2^. Clusters of small diameter axons were identified based on TUJ1 signal using Ilastik 1.0 (Berg et al., 2019). The software was trained to identify clusters of small diameter axons based on morphology, pattern and intensity of TUJ1 signal and ignore single axons or structures that differ from the above. PTBP1 levels were also examined in nuclei of non-neuronal cells based on DAPI positivity.

### Whole tissue staining (iDISCO)

Whole tissue staining was performed as described (Renier et al., 2014) with minor adjustments. Tissue was fixed in 4 % PFA o/n at 4 °C and washed in PBS. Subsequently, samples were dehydrated in methanol/H_2_O series (20 %, 40 %, 60 %, 80 %. 100 % for 1 h each), bleached in fresh 5 % H_2_O_2_ in methanol o/n at 4 °C and rehydrated the next day. After 2 washes in PTx.2 buffer (0.2 % TritonX-100 in PBS) samples were permeabilized in PTx.2 containing 23 mg/ml Glycine and 20 % (v/v) DMSO for 2 d at RT. Blocking was performed in PTx.2 containing 6 % donkey serum and 10 % DMSO for 3 d at RT. ABs for PTBP1, PTBP2 and NFH were applied in in PTwH buffer (0.2 % Tween-20, 0.01 mg/ml Heparin in PBS) supplemented with 5 % DMSO and 3 % donkey serum for 3 d at RT. After 5 washes (each 30 min) in PTwH buffer, tissue samples were incubated with secondary ABs in PTwH with 3 % donkey serum for 2 d at RT, followed 1 h DAPI in PTwH and 5 washes in PTwH (30 min). Finally, samples were transferred to 75 % Glycerol for 1 week before imaging. For imaging, an Olympus FV1000 Confocal laser-scanning microscope at 60x magnification with oil-immersion objective (Olympus UPLSAPO, NA 1.35) was used.

### Fluorescence in-situ hybridization (FISH)

Single molecule fluorescence *in situ* hybridization combined with immunofluorescence (FISH/IF) was used to detect and quantitate *RHOA* and *KPNB1* mRNA and PTBP1 protein in mouse sciatic nerve sections as described (Spillane et al., 2013). 5’ labelled Stellaris probes were designed against mouse *RHOA* and *KPNB1* mRNA. Sciatic nerve segments were fixed overnight in 2% PFA at 4°C and then cryoprotected overnight in 30 % sucrose at 4°C. 25 μm thick cryosections were prepared and stored at −20°C until use. Slides were dried at 37°C for 1 hour and then brought to room temperature; all subsequent steps were performed at room temperature unless indicated otherwise. Sections were washed 10 min in PBS once, then 10 min in 20 mM Glycine three times followed by 5 min in fresh 0.25 M NaBH4 three times. After a 0.1M Triethanolamine (TEA) rinse, sections were incubated in 0.25% Acetic Anhydride in 0.1 M TEA for 10 min, followed by two washes in 2X saline-sodium citrate (SSC) buffer and dehydration in in graded ethanol solutions (70, 95, and 100 % 3 min each). Sections were then delipidated in chloroform for 5 min, rehydrated in graded ethanol (100 and 95 % 3 min each) and equilibrated in 2X SSC. Samples were then incubated at 37°C for 5 min in a humidified chamber in hybridization buffer (10% Dextran Sulfate, 1 mg/ml *E. coli* tRNA, 2 mM Vanadyl Ribonucleosides, 200 ug/ml BSA, 2X SSC, 10% formamide, Roche Blocking buffer) followed by overnight in hybridization buffer with probes (7 μM each), goat anti-PTBP1 (1:100;Abcam ab5642), and RT97 mouse anti-NF (1:100; Dev Hybrid Studies Bank). Sections were washed twice in 2X SSC + 10% formamide at 37°C for 30 min and once in 2X SSC for 5 min. Tissues were permeabilized in PBS + 1% Triton-X100 (PBST) for 5 min, and then incubated for one hour in either Cy3- or Cy5-conjugated donkey anti-goat and FITC-conjugated donkey anti-mouse antibodies(1:200 for each; Jackson ImmunoResearch) in 1X blocking buffer (Roche) plus 0.3 % Triton-X100. After washing with PBS for 5 min, sections were post-fixed in buffered 2% PFA for 15 min, washed in PBS three times for 5 min, rinsed in DEPC-treated water, and mounted using *Prolong Gold Antifade.*

For analyses of FISH signals, images were acquired using Leica SP8X confocal microscope with HyD detectors and Lightening deconvolution. Scramble (SCRM) probe signals were used to set the image acquisition parameters such that all acquisitions were set at the scramble probe parameters that generated least signals. xyz image stacks were obtained using 63x oil-immersion objective (1.4 NA) at two random locations along each nerve section. NIH ImageJ colocalization plug-in (https://imagej.nih.gov/ij/plugins/colocalization.html) was used to extract RNA signals in each optical plane that overlapped with Neurofilament. Quantification of this ‘axon only’ mRNA signal was done by analysis of pixel intensity across each xy plane of the extracted axon only channels using image J. FISH signal intensity was normalized as pixels/μm^2^ of NF signal within each xy plane. The average of FISH signal intensity to NF immunoreactivity in each xy plane was averaged across the image z stack of tile image. The relative mRNA signal intensity was averaged for all tiles in each biological replicate. Using NIH ImageJ plugin JACoP (https://imagej.nih.gov/ij/plugins/track/jacop.html), Pearson’s coefficients for colocalization of axon only PTBP1 protein with either *RHOA* or *KPNB1* mRNA were generated.

### RNA IP and RNA sequencing

Mice were subjected to SN crush and allowed to recover for 7 d before axoplasm collection from injured (inj) and naive (na) nerves, using 20 animals/biological replicate. Before splitting the samples to the experimental conditions (na PTBP1-IP, na Ctrl, inj PTBP1-IP, inj Ctrl) 10 % of naive and injury extract was taken as input sample. IP was performed as described previously (Doron-Mandel et al., 2016). RNA from input and IP samples was extracted using the RNeasy Micro Kit (Qiagen, Cat #: 74004) with on-column DNase treatment according to the manufacturer’s instructions. We performed Western blot on supernatants to validate successful PTBP1-IP (depletion of PTBP1 in supernatants of IP samples, no depletion in Ctrl; Fig. S3A).

RNA sequencing was performed for 3 biological replicates, with 3 experimental conditions (Input, PTBP1-IP, Ctrl) and 2 time points: naive and 7 d after SN crush injury. RNA-sequencing libraries were prepared using the Nugen Ovation RNA Ultra Low Input (500ng) with TrueSeq Nano kit (Illumina). Libraries were indexed and sequenced by HiSeq4000 with 50 bp paired-end reads and at least 59M reads (avg 74.5M) were obtained for each sample.

Quality control was performed on base qualities and nucleotide composition of sequences, mismatch rate, mapping rate to the whole genome, repeats, chromosomes, key transcriptomic regions (exons, introns, UTRs, genes), insert sizes, AT/GC dropout, transcript coverage and GC bias to identify problems in library preparation or sequencing. Reads were aligned to the mouse mm10 reference genome (GRCm38.75) using the STAR spliced read aligner (ver. 2.4.0). Average percentage of uniquely aligned reads were 66.3 %. Total counts of read-fragments aligned to known gene regions within the mouse (mm10) ensembl (GRCm38.p6) transcript reference annotation were used as the basis for quantification of gene expression. Fragment counts were derived using HTSeq program (ver. 0.6.0). Genes with minimum of 5 counts for at least one condition (all replicates) were selected and differentially expressed transcripts were determined by Bioconductor package EdgeR (ver. 3.14.0). Scripts used in the RNA sequencing analyses are available at https://github.com/icnn/RNAseq-PIPELINE.git. RNA seq data are available under GEO # GSE142576.

Canonical pathway analysis was performed using Ingenuity^®^ Pathway Analysis (IPA, version 51963813, QIAGEN Inc., https://www.qiagenbioinformatics.com/products/ingenuitypathway-analysis) (Kramer et al., 2014). A complete list of identified transcripts and IPA analysis can be found in Table S2.

### shRNA mediated knockdown of PTBP1

The design of the shRNA constructs was based on AAV-shRNA-ctrl (Addgene, plasmid #85741) (Yu et al., 2015). shRNA expression is driven from a U6 promoter and the plasmid also contains an EYFP reporter. Adeno associated virus (AAV) serotype 9 was chosen due to its preference for peripheral neurons in adult mice.

shCtrl (5’-GTTCAGATGTGCGGCGAGTGAAGCTTGACTCGCCGCACATCTGAAC-3’) was replaced by

shPTBP1 (5’ -AGCCGCTTTCTGTGCCTTAGAAGCTTGTAAGGCACAGAAAGCGGCT-3’) using BamHI and XbaI restriction sites.

### AAV9 production

Purified AAV9 was produced in HEK 293t cells (ATCC^®^), with the AAVpro^®^ Purification Kit (All Serotypes) from TaKaRa (#6666). For each construct 10 plates (15 cm) were transfected with 20 μg of DNA (AAV-plasmid containing the construct of interest and two AAV9 helper plasmids) using jetPEI^®^ (Polyplus) in DMEM medium without serum or antibiotics. pAAV2/9n and pAdDeltaF6 helper vectors were obtained from the University of Pennsylvania Vector Core. Medium (DMEM, 20 % FBS, 1 mM sodium pyruvate, 100 U/mL penicillin 100 mg/mL streptomycin) was added on the following day to a final concentration of 10% FBS and extraction was done 3 days post transfection. Purification was performed according to the manufacturer’s instructions (TaKaRa, #6666). For both constructs we obtained titers ~ 5×10^14^ viral genomes/ml, which were used undiluted for intrathecal (IT) injections (5 μl/animal) or transduction in culture (1μl/well).

Validation of knockdown efficiency was performed by transduction of DRG neurons in culture for 10 days. Cells were fixed in 4% PFA, stained for PTBP1 and quantified using ImageJ/Fiji (Fig. S4A).

### Knockdown *in vivo*

To knockdown PTBP1 in DRG neurons *in vivo,* we delivered shPTBP1 and shCtrl via intrathecal (IT) injection of AAV9. A total of 5 μl (~ 5×10^14^ viral genomes/ml) was injected using a sterile 10 μl Hamilton micro syringe fitted with a 30-gauge needle. Baseline behavior was assessed 3 weeks after injection and showed no significant differences between groups (Fig. S5). Subsequently, we performed SN crush injury on the right hindleg, the left side was used as an uninjured, naive control. At distinct timepoints after SN crush, mice were subjected to behavioral testing (CatWalk, heat probe, von Frey). A first cohort was sacrificed for tissue collection and DRG culture at day 7, 3-4 mice for each group have been kept for longer time (28 days) to monitor functional recovery.

### Conditioning lesion culture

For growth assays after conditioning lesion (Smith and Skene, 1997), L4 & L5 DRG were extracted separately from both sides (left side: naive; right side: 7d after SN crush) from both experimental groups. Cells were allowed to grow for 20 h after plating. Imaging and analysis were performed using ImageXpress Micro (Molecular Devices) automated microscopy system with MetaXpress analysis software (Molecular Device, Version 5.1). Cells, with the longest neurite > 2x the diameter of the cell body, were considered as “growing”; and only EYFP positive neurons have been included in the outgrowth analysis.

### Sciatic nerve longitudinal sections

Tissues were fixed in 4 % PFA for 24h at 4 °C, then washed in PBS, followed by incubation in 30 % sucrose for 48 hours at 4 °C. Tissues were embedded in O.C.T. Compound (Scigen, 4583) and 15 μm sections were obtained. Blocking was performed in 10 % horse serum, 0.2 % Triton X100 in PBS for 2 h at RT. Primary antibodies were incubated in AB-solution (5% horse serum, 0.2 % Triton X-100 in PBS) overnight at 4 °C. Secondary ABs were applied in AB-solution for 2 h at RT. Slides were mounted in Fluoromount (F4680, Sigma Aldrich). Samples were acquired using a Panoramic MIDI slide scanner (www.3dhistech.com) equipped with a 20X objective. For the analysis, images were taken in a range between from 1 to 4 mm proximal and distal from the injury site. For each image, CGRP or NFH positive fibers were counted in three regions of interest (size 40 microns each) orthogonal to the fiber direction, in the center and on the sides of the image. The number of fibers was normalized by the diameter of the nerve section.

### PTBP1 inhibition via decoy RNA oligo

Inhibitory decoy RNA oligonucleotides for PTBP1 (PTBP1i) and scramble control (SCRMi) were as previously described (Denichenko et al., 2019). These oligonucleotides are single stranded RNA molecules of three/four tandem motif repeats with a 2’-O-methyl modification on the ribose of each nucleotide. To increase stability for *in vivo* experiments, we modified the oligos with a phosphorothioate backbone and also added a fluorophore (TYE665) to the 5’ end to allow for imaging. Oligos were synthesized by IDT (Integrated DNA Technologies, Syntezza, Israel).

SCRMi: 5’TYE665-GCAAUCCGCAAUCCGCAAUCC-3’

PTBP1i: 5’TYE665-CUCUCUCUCUCUCUCUCUCUCUCU-3’

Decoy RNA oligos were used to locally inhibit PTBP1 in the SN and to verify its role after SN crush injury. Therefore, we injected 865ng of PTBP1i or SCRMi oligo (in a volume of 2 μl) directly into the SN using a sterile 10 μl Hamilton micro syringe fitted with a 34-gauge needle. Immediately after, we crushed the sciatic nerve in the same position. The other side served as a naive control. Mice were subjected to behavioral testing (CatWalk, heat probe, von Frey) in a similar manner as in the knockdown experiment.

### Statistical analysis

All analyses were performed using GraphPad Prism version 8.43 for Windows (GraphPad Software, La Jolla, California, USA, www.graphpad.com). Statistically significant p values are shown as * p < 0.05, ** p < 0.01, *** p < 0.001 and **** p < 0.0001. All data underwent normality testing using the Shapiro-Wilk test. Datasets that passed the normality test were subjected to parametric analysis. Unpaired Student’s t-test was used for analysis with 2 groups, one-way ANOVA was used to compare multiple groups, two-way ANOVA or Mixed-effects analysis was used to compare 2 variables (PTBP1 perturbation, SN injury). Paired t-test or repeated measure (RM) analysis was done on data coming from the same animal. Tukey’s, Sidak’s or Dunnett’s multiple comparisons tests were used in the follow up analysis as specified in the figure legends, usually comparing experimental conditions to control (naive). Data sets that did not pass the normality test were subjected to nonparametric analysis using Mann-Whitney test for 2 groups, or Kruskal-Wallis test for multiple group evaluation, followed by Dunn’s multiple comparisons test. The results are shown as mean ± standard error of the mean (SEM). All statistical parameters for each analysis are stated in the corresponding figure legends.

---

**Fig. S1:**
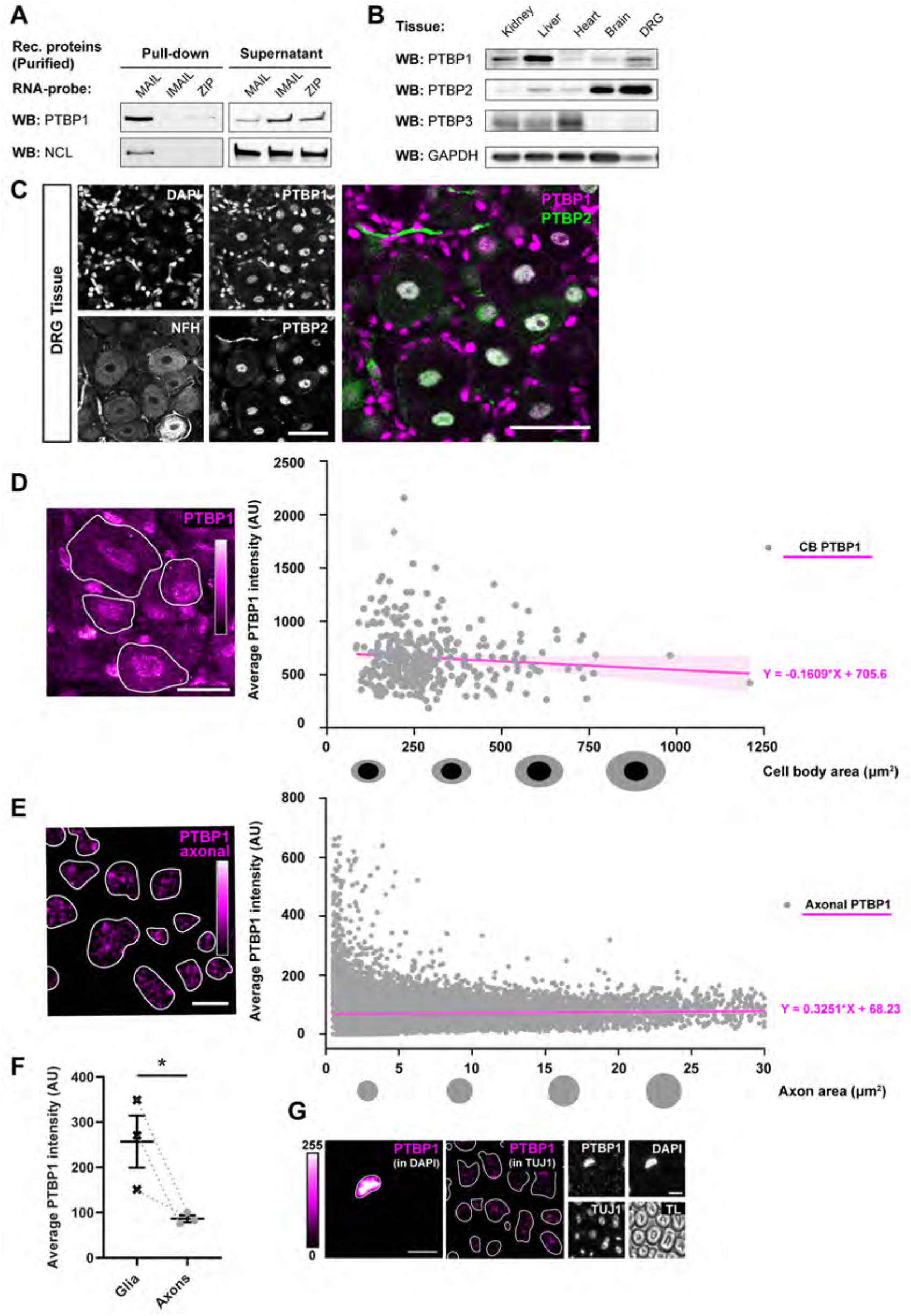
PTBP1 is expressed in adult sensory neurons and associated with *KPNB1* mRNA. (**A**) *In vitro* binding assay using recombinant PTBP1 or Nucleolin (NCL) with MAIL (motif for axonal importin localization), IMAIL (inactive MAIL) or zipcode motif (ZIP) of β-Actin. Detection by western blot (WB). This assay proves a direct association of PTBP1 with the RNA localization motif of *KPNB1.* (**B**) WB for different PTBP paralogues (PTBP1, PTBP2, PTBP3) in several tissues. GAPDH as housekeeping gene, n = 6 mice for PTBP1 in liver, brain and dorsal root ganglia (DRG), quantified in Fig. 1D. (**C**) Whole tissue staining (iDISCO) on lumbar DRG: co-localization of PTBP1 and PTBP2 in the nucleus of DRG neurons. PTBP2 strictly neuronal, PTBP1 expression in neurons and glia cells. n = 3 mice, scale bar = 50 μm, NFH = Neurofilament heavy chain. (**D**) PTBP1 expression in DRG neurons of all sizes. Simple linear regression for PTBP1 intensity in the cell body (CB) versus cell body area (μm^2^), n = 330 neurons (from 4 mice), regression line ± 95% CI in magenta, R^2^ = 0.008524. **e**, PTBP1 levels in axons of the sciatic nerve. Simple linear regression for axonal PTBP1 versus axon area (μm^2^), n = 24,736 axons (from 3 mice), regression line ± 95% CI in magenta, R^2^ = 0.0009398. Objects >30μm^2^ have been discarded for the analysis, for illustration purposes 6 datapoints with PTBP1 intensity >800 are not shown (but included in liner regression analysis). **D & E** show that in DRG neurons PTBP1 is expressed in neurons of different CB sizes and axons of different diameters. Images in **D** & **E** are not exposure matched (independent experiments, different tissues). They show PTBP1 in Magenta HOT scale (0-255), outline of TUJ1 indicated with white dashed line, scale bar **D** = 20 μm, scale bar **E** = 5 μm. (**F**) Quantification of PTBP1 in glial nuclei vs. TUJ1 positive axons within the same image. Individual datapoints with mean ± SEM, ratio paired t-test, p-value * < 0.05, n = 3 mice (> 1000 axons for each mouse). Grey dashed line indicates paired datapoints. (**G**) Images corresponding to **F**, with panels for TL (transmitted light), DAPI, TUJ1 and PTBP1. Nuclear and axonal signal for PTBP1 is shown in magenta hot scale (exposure matched, same image, 0-255), with the mask for DAPI or TUJ1 as a white outline. Scale bar = 5 μm.

**Fig. S2:**
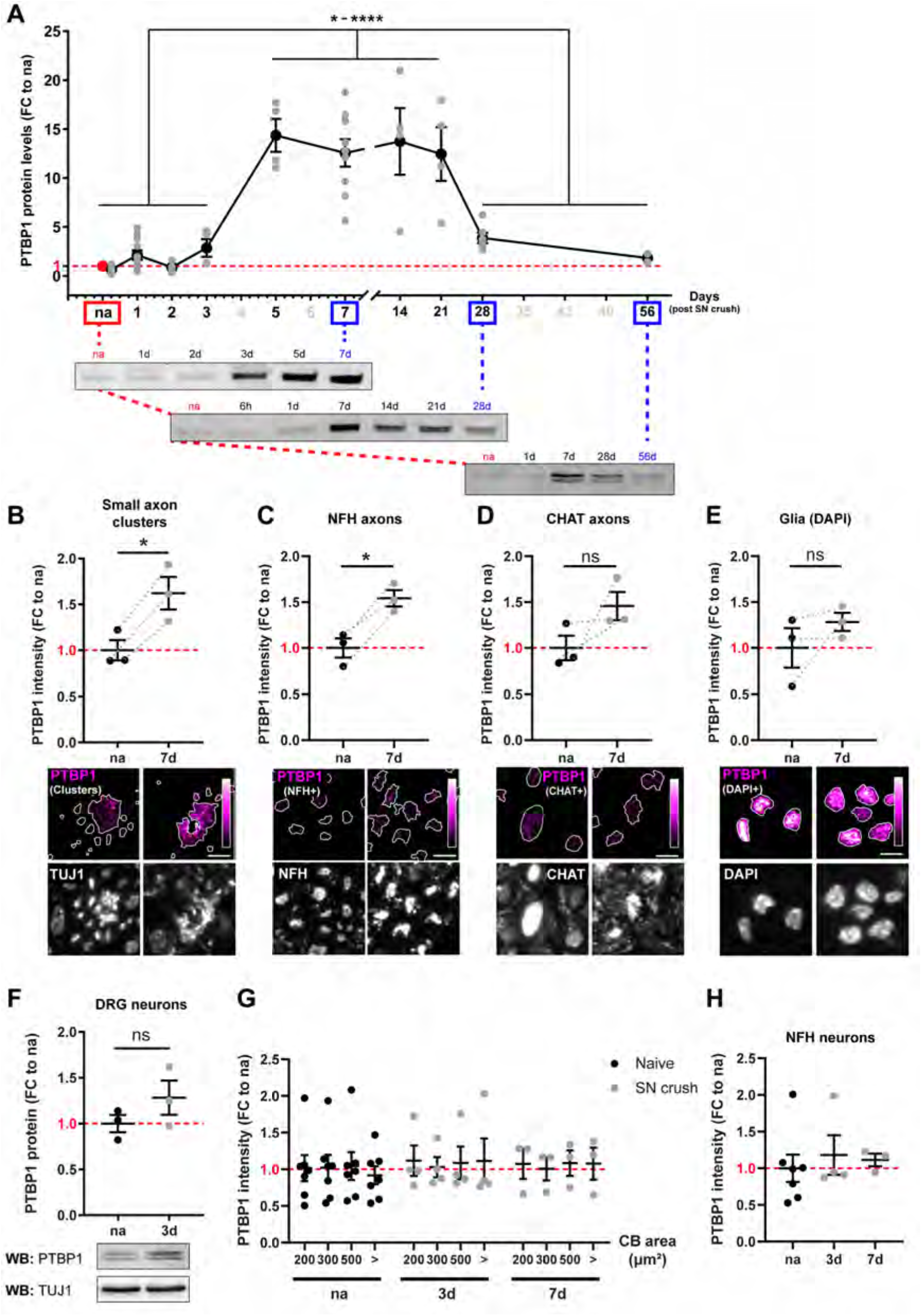
Increased levels of PTBP1 in axons 7d after SN crush. (**A**) Quantification of PTBP1 levels in sciatic nerve (SN) axoplasm under naive (na) and injury (inj) conditions, combining several experiments. Detection by western blot (WB). PTBP1 levels were normalized to a housekeeping gene and the data is shown as fold change (FC) to naive, which was used in all experiments as a reference (red dashed line). The data is shown as mean ± SEM (black curve) with an overlay of the individual data points in grey. One-way ANOVA with Tukey’s multiple comparison test. Time points 5-21 d are significantly different compared to na, 1d, 2d, 3d, 28d and 56d. p-value * -**** < 0.05-0.0001, n = 4-11 mice for each timepoint. Most relevant timepoints for this study are shown also in Fig. 2A (Experimental set with 3 replicates, performed with the same blotting conditions). **B-E**, Quantification of PTBP1 intensity in different categories of axons and glia nuclei. Individual datapoints with mean ± SEM, ratio paired t-test, p-value * < 0.05, n = 3 mice (>1000 positive objects/mouse). (**B**) Clusters of small axons (TUJ1, Tubulin beta class III); (**C)** NFH (Neurofilament heavy chain) positive axons; (**D**) CHAT (Choline O-Acetyltransferase) positive axons; (**E**) Glia cell nuclei (DAPI positive objects). Images below show PTBP1 in magenta HOT scale within the outlines of the corresponding mask and immunofluorescent images for the corresponding markers, scale bar = 5 μm. (**F**) PTBP1 protein expression in purified neurons of DRG tissue (cells after percoll gradient). WB of PTBP1 with normalization to TUJ1, shown as FC to naive (red dashed line). Individual datapoints with mean ± SEM, t-test, not significant, n = 3 mice. (**G**) Quantification of PTBP1 signal in DRG neurons according to different cell body sizes (Binning of cell body (CB) area: 0-200, 200-300, 300-500, >500 μm^2^). PTBP1 intensity is shown as FC to naive (red dashed line), individual datapoints with mean ± SEM, two-way RM ANOVA with Tukey’s multiple comparison test, not significant, n = 3-7 mice. (**H**) Quantification of PTBP1 signal in NFH positive neurons. FC to na (red dashed line), individual datapoints with mean ± SEM, one-way ANOVA with Tukey’s multiple comparison test, not significant, n = 3-7 mice.

**Fig. S3:**
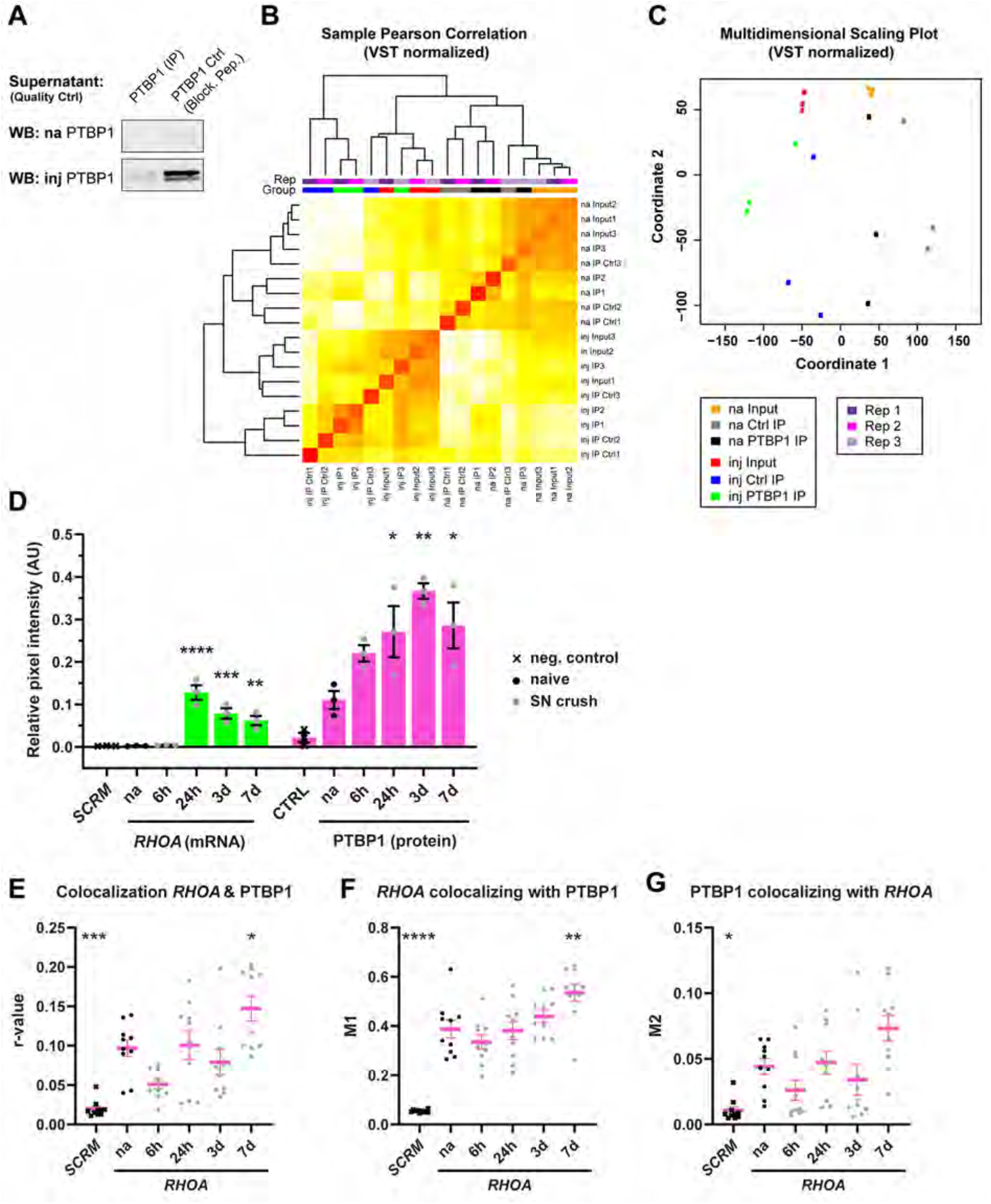
Identification of PTBP1 cargo mRNAs in the SN. (**A**) Quality control for PTBP1 protein IP. Western blot (WB) of supernatants shows depletion in the IP condition and not in the Ctrl (blocking peptide). Enrichment of PTBP1 in injury (inj) condition compared to naive (na). (**B**) Sample Pearson’s correlation including dendrogram. Raw counts were normalized by variance stabilized transformation (VST) after filtering transcripts with low counts. (**C**) Multidimensional scaling plot showing clustering of the samples. **b-c**, Top 5000 genes (according to rank) have been used in the graphs; scripts used in the RNA sequencing analyses are available at https://github.com/icnn/RNAseq-PIPELINE.git. RNA seq data are available under GEO # GSE142576. **D-G**, Extended time course for the RNA-FISH experiment. Individual datapoints with mean ± SEM, one-way ANOVA with Dunnett’s multiple comparison test against naive (na), p * < 0.05, ** < 0.01, *** < 0.001, **** < 0.0001, n = 10 nerve sections from 3 mice. (**D**) Total amount of *RHOA* mRNA and PTBP1 protein signals summarized for each mouse. Separate quantification for mRNA and protein signal. (**E**) Pearson’s correlation analysis of axonal PTBP1 with *RHOA*, r-value. (**F**) M1. (**G**) M2.

**Fig. S4:**
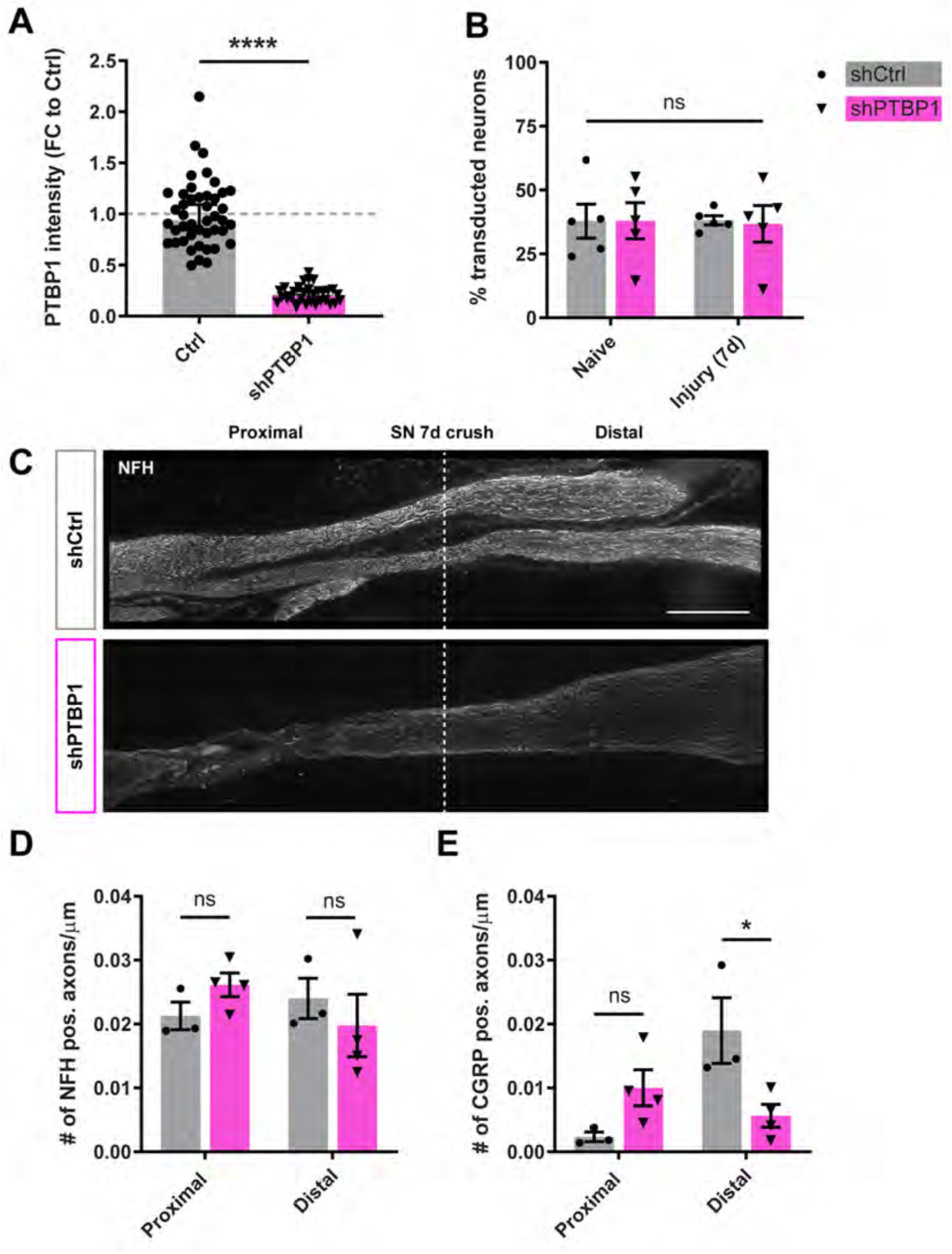
PTBP1 plays a role in the injury response of adult sensory neurons. (**A**) Validation of shRNA in DRG culture. Transduction *in vitro* and culture for 10 days, positive cells (cells transduced with shPTBP1) were compared to negative control cells (Ctrl). Quantification of PTBP1 protein levels by immunofluorescence (IF) in EYFP positive cells vs. EYFP negative cells from the same mouse. Data is shown as FC (fold change) to Ctrl with individual datapoints and mean ± SEM, Mann-Whitney test, p-value **** < 0.0001, n > 30 cells. (**B**) Validation of transduction efficiency *in vivo*. % of positive neurons was determined by IF EYFP expression) 4 weeks after intrathecal injection of AAV9-shRNA (3 weeks +7 days crush). DRG have been dissociated for culture, fixed 20 h after plating, and stained for TUJ1 and EYFP. Transduction ~ 37% with no difference between shRNA constructs or left (naive) and right (injury) site. Individual datapoints with mean ± SEM, two-way repeated measure (RM) ANOVA with Sidak’s multiple comparison test, n = 5 mice. (**C**) SN longitudinal sections 7d after crush for shCtrl and shPTBP1. Staining for NFH (Neurofilament heavy chain), injury site indicated by the white dashed line, scale bar = 500 μm. **D** & **E,** Quantification of the number of positive axons (average of several regions up to 4 mm) proximal and distal to the crush site. Individual datapoints with mean ± SEM, two-way RM ANOVA with Sidak’s multiple comparison test, n = 3-4 mice. (**D**) NFH positive axons, not significant (ns). (**E**) CGRP (Calcitonin Gene-Related Peptide 1) positive axons, p-value * < 0.05.

**Fig. S5:**
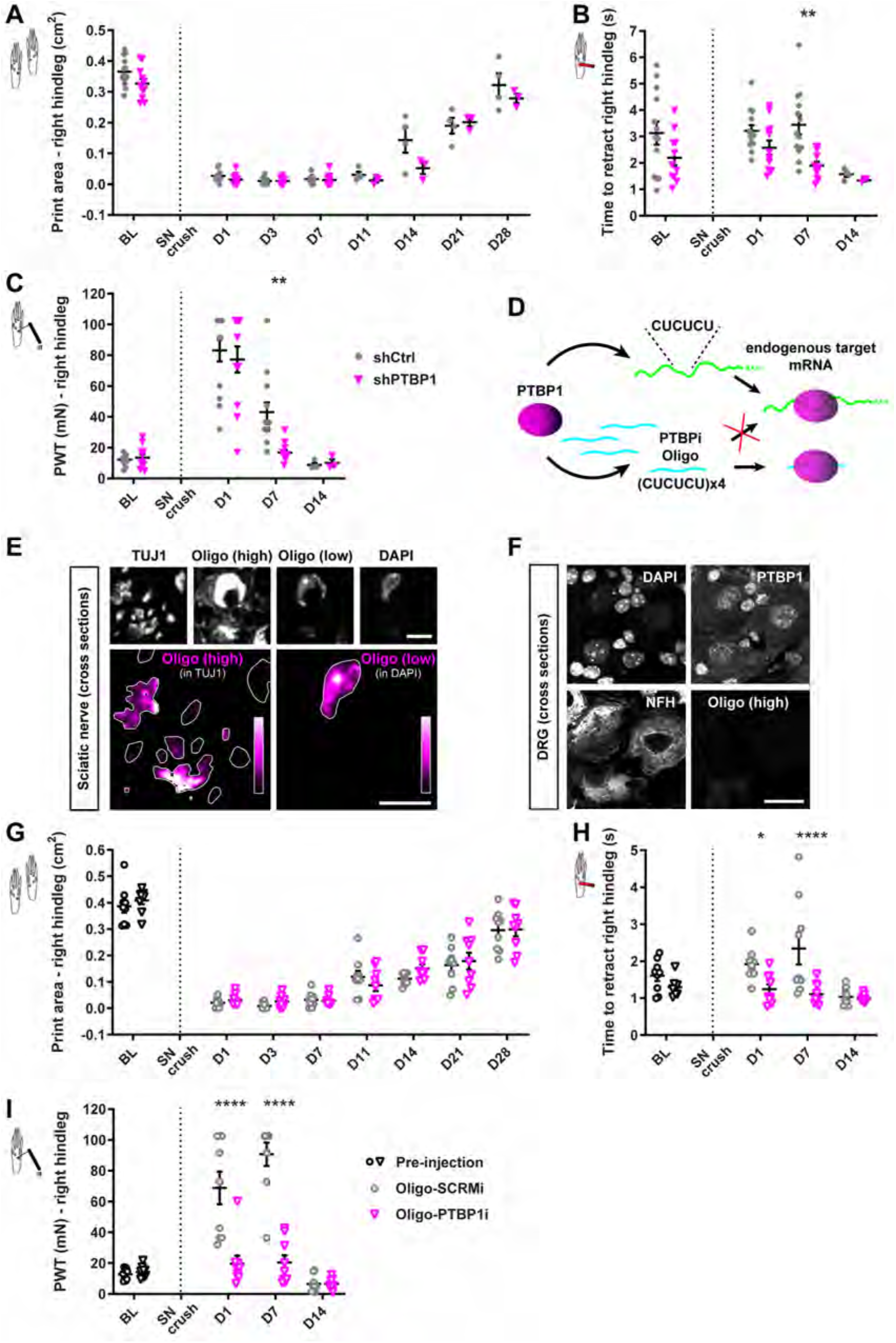
PTBP1 perturbation after SN crush leads to altered sensation in vivo. **A-C,** Behavior, after knockdown of PTBP1, following sciatic nerve (SN) crush injury. Baseline (BL) testing was performed 3 weeks after AAV injection of shRNA, just prior to crush. Mice have been tested in CatWalk gait analysis **(A)**, heat probe **(B)** and von Frey **(C)** paradigms. PWT (paw withdrawal threshold). Individual datapoints with mean ± SEM. BL to day 7: n = 11-13 mice; day 8 onwards: n = 3-4 mice. Mixed-effects analysis with repeated measures (RM) followed by Sidak’s multiple comparison test, p-value ** < 0.01 for heat probe and von Frey, CatWalk gait analysis is not significant. Analysis for both hindlimbs left (naive), right (SN crush) at day 7 is shown in **Fig. 5 B-D**. (**D**) Schematic showing the mechanism of the decoy RNA oligo (Denichenko et al., 2019). The RNA decoy oligo for PTBP1 (PTBP1i, cyan) consists of the repetitive CU-binding motif and prevents PTBP1 protein (magenta) from binding to its endogenous mRNA targets (green). For *in vivo* experiments, decoy RNA oligos were labeled with a TYE665 fluorophore on the 5’ end for imaging and were modified with a phosphorothioate backbone and 2’-O-methyl modifications. **e**, Images of sciatic nerve (SN) cross sections after oligo injection with SN crush. Oligo (TYE665 label) was detected with high and low laser intensities to show colocalization with TUJ1 or DAPI, respectively. Big images below show oligo intensity in Magenta HOT scale (0-255) within TUJ1 or DAPI positive objects (white outline). Images are overexposed for illustration purposes, scale bar = 5 μm. (**F**) Images of dorsal root ganglia (DRG) cross sections after oligo injection into the SN. No signal for the oligo was detected in the cell bodies of DRG neurons. Scale bar = 20 μm. **G-I,** Behavior, after inhibition of PTBP1 via decoy RNA oligo, following sciatic nerve (SN) crush injury. BL testing was performed just prior to injections and SN crush. Mice have been tested in CatWalk gait analysis **(G)**, heat probe **(H)** and von Frey **(I)** paradigms. Individual datapoints with mean ± SEM, n = 9 mice. Two-way RM ANOVA with Sidak’s multiple comparison test, p-value * < 0.05, **** < 0.0001 for heat probe and von Frey, CatWalk gait analysis is not significant. Analysis for both hindlimbs left (naive), right (SN crush) at day 7 is shown in **Fig. 5 F-H**.

